# Milti-Scale Detrended Partial Cross-Correlation Analysis of Tree Ring Width and Climate Variations: Revealing Heat and Drought Stress Resilience Factors in a Forest Ecosystem

**DOI:** 10.1101/2023.05.30.542825

**Authors:** Mikhail I. Bogachev, Artur M. Gafurov, Pavel Y. Iskandirov, Dmitrii I. Kaplun, Airat R. Kayumov, Asya I. Lyanova, Nikita S. Pyko, Svetlana A. Pyko, Anastasia N. Safonova, Aleksandr M. Sinitca, Bulat M. Usmanov, Denis V. Tishin

**Affiliations:** St. Petersburg Electrotechnical University “LETI”, 5 Professor Popov street, St. Petersburg, 197022, Russia; Kazan Federal University, 18 Kremlevskaya street, Kazan, 420008, Tatarstan, Russia

**Keywords:** Scots pine, tree-ring width, drought stress, hydrological conditions, detrended partial cross-correlation analysis

## Abstract

In a changing climate, forest ecosystems become increasingly vulnerable to the continuously exacerbating heat and drought stress conditions. Climate stress resilience is governed by a complex interplay of global, regional and local factors, with hydrological conditions among the key roles. Using a modified detrended partial cross-correlation analysis (DPCCA), we analyse the interconnections between long-term tree-ring width (TRW) data and regional climate variations at various scales and time lags. By comparing dendrochronological series of Scots pine trees near the southern edge of the boreal ecotone, we investigate how local hydrological conditions affect heat- and drought stress resilience of the forest ecosystem. While TRW are negatively correlated with spring and summer temperatures and positively cor-related with the Palmer drought severety index (PDSI) in the same year indicating that heatwaves and droughts represent the limiting factors, at interannual scales remarkable contrasts can be observed between areas with different local hydrological conditions. In particular, for the sphagnum bog area positive TRW trends over several consecutive years tend to follow negative PDSI trends and positive spring and summer temperature trends of the same duration with a time lag between one and three years, indicating that prolonged dry periods, as well as warmer springs and summers appear beneficial for the increased annual growth. In contrast, for the surrounding elevated dry land area a reversed tendency can be observed, with pronounced negative long-term correlations with temperature and positive correlations with PDSI. Moreover, by combining detrending models and partial correlation analysis, we show expicitly that the long-term temperature dependence could be partially attributed to the spurious correlations induced by coinciding trends of the trees ageing and climate warming, while contrasts in correlations between TRW and PDSI become only further highlighted, indicating the major impact of the local hydrological conditions on the drought stress resilience.

**Graphical Abstract:** 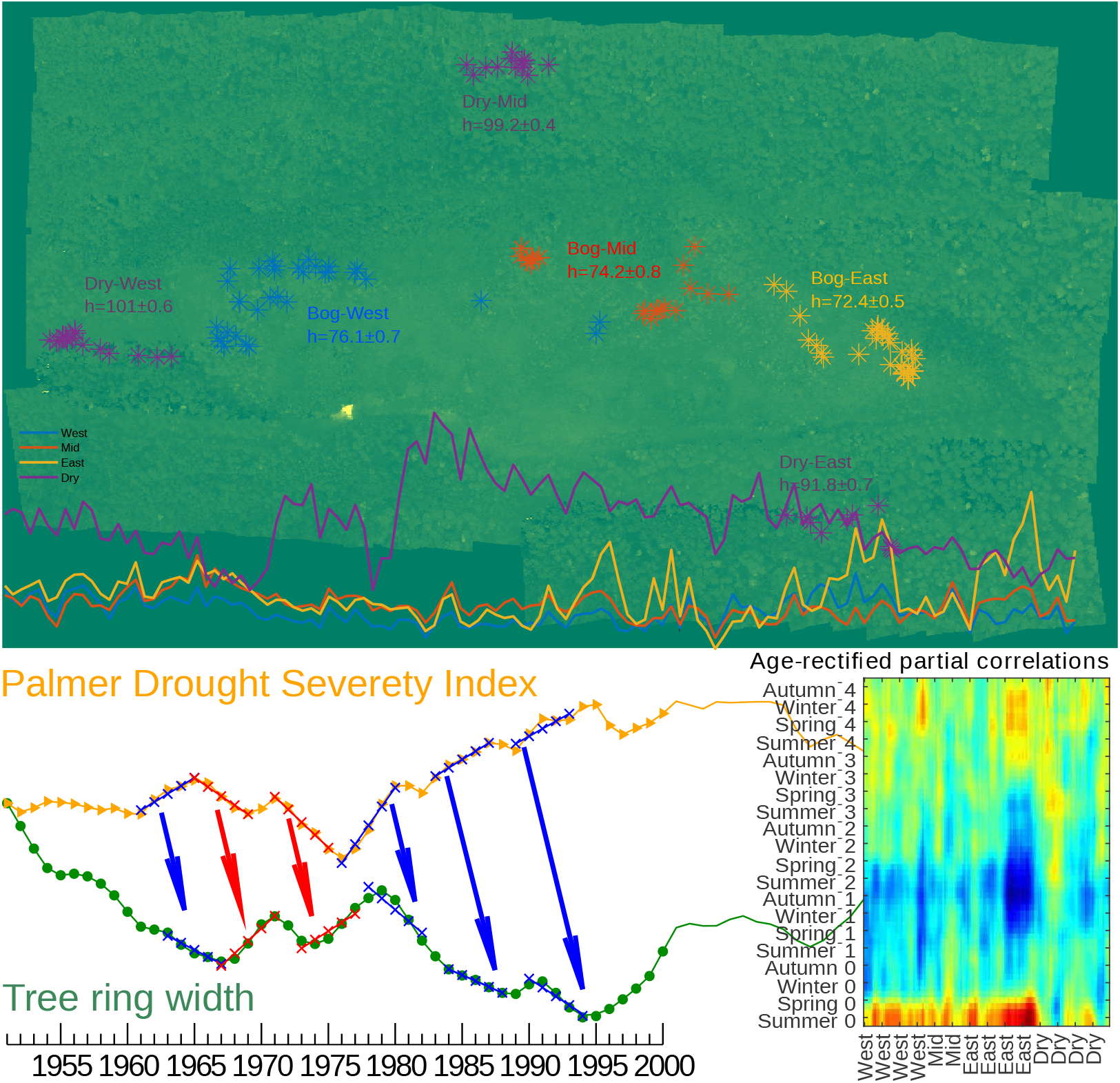

**Highlights:**

- Climate stress resilience of forest ecosystems is largely driven by local hydrology
- Multiscale analysis reveals reversed climate stress response in dry and wetland areas
- Warmer springs and summers are favorable for tree growth under nondrought conditions
- Warm and dry periods improve trees growth in the peat bog area with 1-3 year time lag
- Climate stress response and ageing effects can be understood from partial correlation

## 1. Introduction

In a changing climate, the duration of summer heat waves in Europe has doubled, and the number of days during which extreme temperatures have been recorded has tripled since 1880 [1]. Among recent examples, in addition to intensified flash droughts [2], the 2010 extreme heatwave accompanied by a prolonged drought [3] led to increased demand for evaporation [4], exacerbating the negative impact of extreme temperatures on the ecosystems productivity, vigor, and survival [5].

Forest ecosystems play an increasingly indispensable role in the preservation of the microclimate conditions essential to the balance of the environment and the well-being of the local biome as a whole. Climate exerts the strongest control over the geographic location of ecotones, with coniferous forests located near the southern edge of the boreal ecotone being among the most endangered due to the increased environmental pressure. While recent shifts of the ecotone boundaries to higher latitudes under climate change could hardly be counteracted, with certain alterations seemingly irreversible at the continental scale, fine-scale drivers play an increasingly important role in the definition of the biome boundaries, and thus also in the preservation and well-being of the local ecotones [6].

Among prominent examples, conifer trees of the *Pinaceae* family are very sensitive to summer droughts, and thus coniferous forests growing near the southern border of the boreal zone could be also considered as characteristic bioindicators highly sensitive to hydroclimatic changes and overall environmental pressure. Under combined drought and heat stress, a decrease in photosynthesis activity significantly limits carbon uptake by the ecosystem. As soils dry out and tree canopy transpiration exceeds water uptake by roots, tree stem water is gradually depleted. The depletion of stem water stunts tree growth, further reducing the capacity of forests to absorb carbon. In the short term, the release of water from the internal reserves of the trunk can temporarily mitigate the negative impact of drought on the integrity of the tree’s vascular system. However, extended periods of drought eventually lead to hydraulic failure as well as dehydration and tissue damage, which can lead to tree death due to drought [7].

Accordingly, unraveling the complex interplay of global, regional and local factors governing heat and drought stress resilience of the endangered forest ecosystems is of immense importance for a better understanding of their adaptability in a changing climate, and thus also for the improvement of the local environmental management. Long-term persistence in climate variability [8] leads to both clustering of flash heatwaves and emergence of prolonged droughts [9], this way increasing the impact of hydroclimatic anomalies on the forest ecosystems considerably. Conventional time series analysis methods often neglect long-term memory, resulting, for example, in the underestimation of clustering effects in tree-ring based climate reconstructions [10] and overestimation of the significance of local trends [11, 12].

In recent years, significant progress have been made in overcoming the above limitations. Long-term memory in climate variability records, their tree-ring data based reconstructions, and surrogate data generated by climate models have been quantified using dedicated methods capable of adequately assessing long-term persistence in the presence of additive short-term and/or periodic trends [13] such as detrended fluctuation analysis (DFA) [14] and wavelet transform analysis (WTA) [15]. Based on the results of these analyses, more accurate methods to evaluate statistical significance of local trends [16, 17, 18], as well as corresponding corrections to tree-ring based climate variability reconstruction procedures [19, 20, 21] have been proposed.

In this work, we extend the detrended fluctuation analysis methodology to the interconnections between hydroclimatic variations and tree-ring width data by substituting the conventional cross-correlation analysis by the corresponding generalizations of detrending methods. Besides the recently proposed Detrended (Partial) Cross-Correlation Analysis (D(P)CCA) [22, 23], we also considered a similar generalization of the central moving average (CMA) based detrended fluctuation analysis [24] which has been proposed to minimize discontinuity in the residuals. Finally, to analyze possible delayed response of the trees growth rates to hydroclimatic variations, we analyzed not only synchronous, but also delayed cross-correlations with time lages up to five years.

## 2. Materials and Methods

### 2.1. Study area

The study area is located in the Volzhsko-Kamsky State Nature Biosphere Reserve on the left bank of the Volga river, 30 km west of Kazan city, Republic of Tatarstan, Russia, near the southern edge of the boreal ecotone. The landscape exhibits a dune-hilly character, with hollows and ancient beams, with absolute heights between 65 and 105 m. Soils are sandy, medium podzolic. Peat bogs of limnogenic origin are typically located in the interdune depressions. The main species of the Volga-Kama Reserve is Scots pine (*Pinus sylvestris* L.). It accounts for up to 70% of the forested area, being represented in various habitats (from bog to dry land).

Our studies focused on the local area of the “Dolgoe” sphagnum bog (N 55.900190, E 48.821186), characterized by the peat thickness up to 3.5 m, and a surrounding elevated dry land area (considered as a relevant control). The local forest type is dominated by sphagnum pines with only sparsely represented spruces and birches. The grass layer is represented by *Oxycoccus palustris*, *Chamaedaphne calyculata*, *Rhododendron tomentosum, Andromeda polifolia* L. and moss *Sphagnum* sp. The studied area shown in Fig. 1 is characterized by a local height gradient leading to a respective gradient in the local hydrological conditions both between the elevated dry land surronding the peat bog and the peat bog area itself, as well as a less pronounced lowering from the western part towards the eastern part of the bog area.

**Figure 1:**
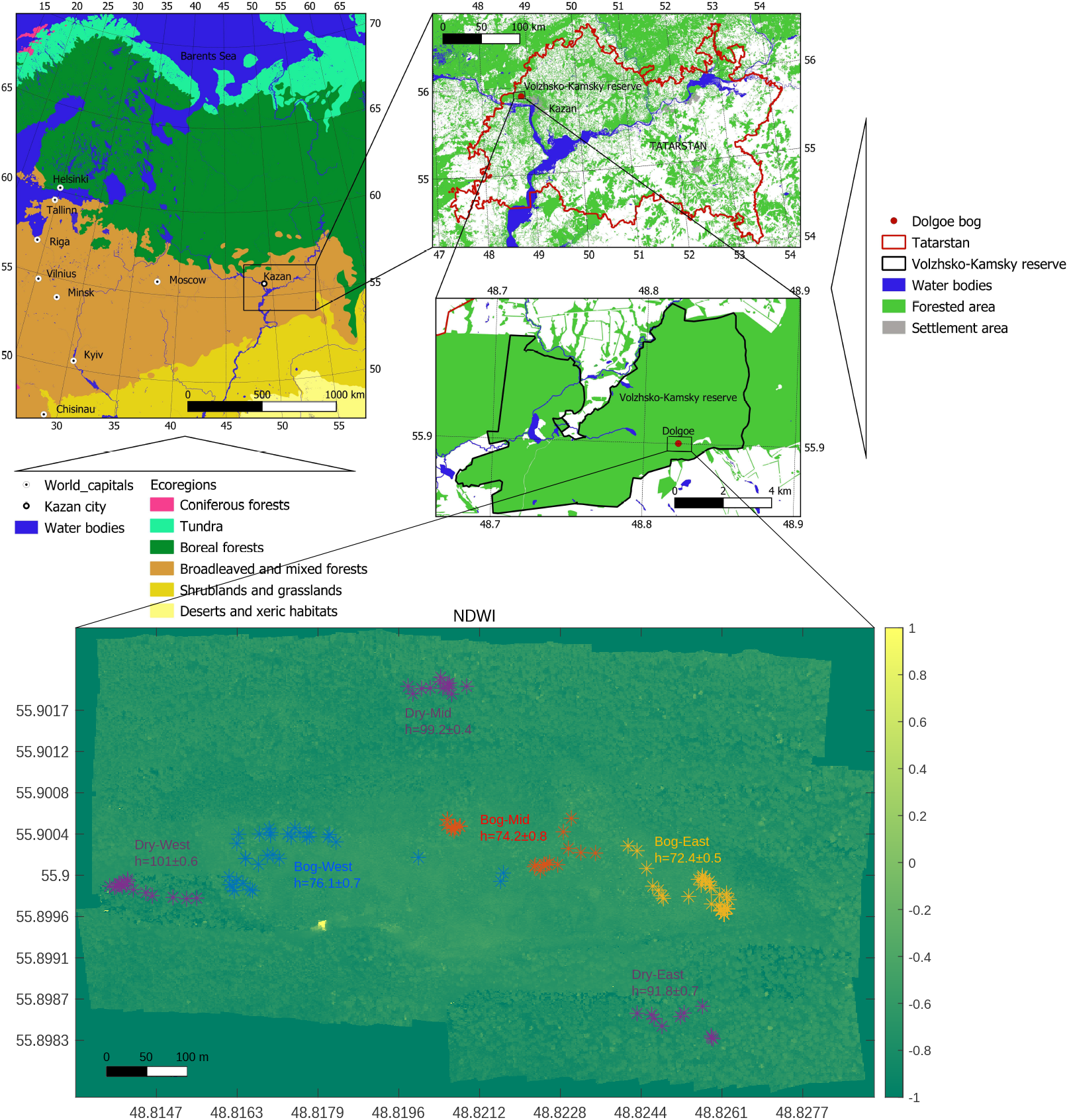
Overview of the studied area with marked up locations of the trees. The geographical location in the upper panel has been prepared using QGIS software based on OpenStreetMap data. Ecoregions have been classified according to [25]. The background in the lower panel represents the NDWI index (for details, see Sec. 2.4) obtained by multispectral imaging.

### 2.2. Tree-Ring Measurements

Trees sampling was carried out according to the methodology adopted in earlier dendrochronological studies [26]. The cores were extracted from *n* = 140 trees growing in different parts of the bog (*n* = 32 in the western, *n* = 26 in the central, *n* = 42 in the eastern), as well as in the surrounding elevated dry land area (*n* = 40 altogether in three surrounding locations, as indicated in Fig. 1) with a Pressler borer. Tree ring width (TRW) was measured on a Lintab-6 with the TSAPWin software package [27]. The quality of the cross-chronologies was assessed using the Cofecha software [28]. The exact location of each studied tree was determined using a Garmin GPSMAR 62S GPS receiver as indicated in Fig. 1.

### 2.3. Meteorological Data Sources

Local temperature variations have been obtained from the RIHMI-WDC Baseline Climatological Database http:\\meteo.ru\english\data [29], Palmer Drought Severity Index (PDSI) data have been obtained from the KNMI Climate Explorer available online at https://climexp.knmi.nl/ [30, 31], both with monthly resolution since 1901.

### 2.4. Multispectal Remote Sensing

Remote sensing imagery has been acquired using Geoscan 401 Geodesy UAV equipped with a MicaSense RedEdge-MX (RX02 series) multi-spectral camera (pixel size 3.75*µm*, resolution 1280 × 960 (1.2 MP ×5 imagers), aspect ratio 4:3, sensor size 4.8 mm × 3.6 mm, focal length 5.4 mm, field of view: 47.2 degrees horizontal, 35.4 degrees vertical, output depth 12 bit). The recorded channels are outlined in the Table 1.

**Table 1:** Spectral channels used in remote sensing analysis.

| Channel | Center ± bandwidth, nm |
| --- | --- |
| BLUE | $475 \pm 32$ |
| GREEN | $560 \pm 27$ |
| RED | $668 \pm 16$ |
| Red edge (RE) | $717 \pm 12$ |
| Near infrared (NIR) | $842 \pm 57$ |

Based on the multispectral remote observations data, the following vegetation indices have been calculated (for their more detailed description, we refer to [32, 33, 34, 35, 36], see also [37, 38, 39] and references therein):

- **Enhanced Vegetation Index (EVI)**

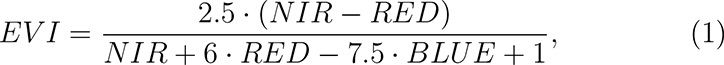 a modification of the conventional Normalized Difference Vegetation Index (NDVI) index that effectively displays greenness (associated with the relative biomass). While the conventional NDVI index takes advantage of the contrast of the characteristics of the chlorophyll pigment absorptions in the red band and the high reflectivity of plant materials in the NIR band, the EVI additionally accounts for atmospheric influences and vegetation background signal, making it less sensitive to background and atmospheric noise, as well as reducing saturation for areas with dense green vegetation.
- **Normalized difference water index**

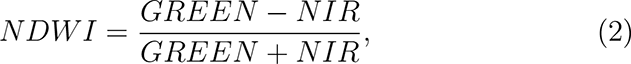 an index originally suggested in [40] to quantify water content, that is simply reciprocal to Green Normalized Difference Vegetation Index (GNDVI), another widely adopted vegetation index that determines water and nitrogen uptake into the plant canopy.
- **Red-Edge Triangulated Vegetation Index (RTVICore)**

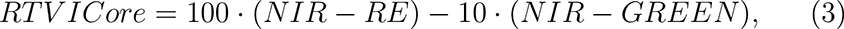 a three-band vegetation index used for estimating leaf area index and biomass. This index uses reflectance in the NIR, red-edge, and green spectral bands.
- **Modified Triangular Vegetation Index (MTVI2)**

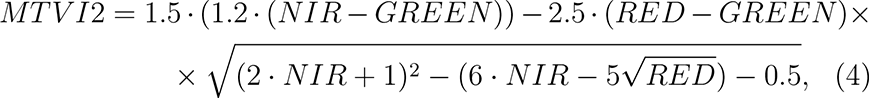 a vegetation index for detecting leaf chlorophyll content at the canopy scale while being relatively insensitive to leaf area index. It uses reflectance in the green, red, and NIR bands.

In this work, we used the NDWI index calculated as indicated in Eq. 2 [40] to quantify the local variations of the relative water content in the soil over the study area, since conventional methods to measure water content in the soil moisture could not be applied in the peat bog area due to saturation. In addition, the EVI, RTVICore, and MTVI2 indices reflect the status of vegetation at the time of observation (17 August 2022).

### 2.5. Time Series Analysis

#### 2.5.1. Detrended Fluctuation Analysis

To quantify long-term persistence in the annual tree-ring data and hydrometeorological variations, we employed the modifications of widely adopted detrended fluctuation analysis (DFA) methods [14]. Since we are particularly interested in a detailed analysis of the impact of short-term vs long-term droughts, as well as flash droughts associated with heatwaves, we take the advantage of the multi-scale fluctuation analysis to extract the relevant information from the observational data.

Technically, multi-scale fluctuation analysis does not deal directly with the observational data series *x_i_*, but with their cumulative sums 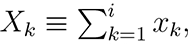 where *i* = 1*, . . ., L* is the data sample, and *L* is the length of the data, also known as “profiles”. In the context of TRW series, the profile represents the cumulative tree growth over its entire lifespan. In order to focus on the relative effects characterized by the TRW fluctuations, without loss of generality, one can subtract the average 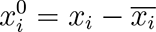 from the raw data series and calculate of the profile for the 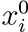 instead of *x_i_*, such that upward and downward trends indicate increasing and decreasing growth rates, respectively.

Next, the profiles are split into *K_s_* time windows of length *s*. In the conventional fluctuation analysis (FA), one considers the mean square displacements over all windows of size *s*

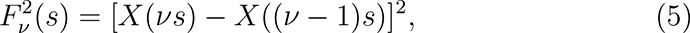

where *ν* is the window number, and averages over all *K_s_* segments for each window size *s* to obtain the fluctuation function *F* (*s*),

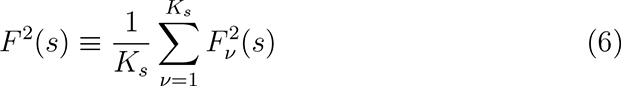

In the DFA procedure, in each window the least mean squares polynomial fits *p_ν,i_* to the profile is calculated, representing local trends in the growth over *s* years. By subtracting the polynomial fits *p_ν,i_* one obtains the fluctuations around the profile and calculates statistics to (5) and (6) as

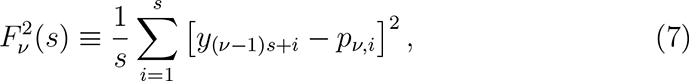

followed by averaging 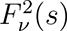 over all *K_s_* windows

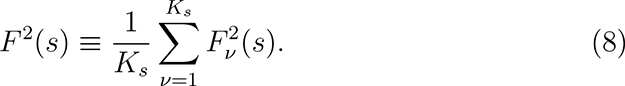

It is known that for long-term correlated data the DFA fluctuation function increases by a power law *F* (*s*) ∝ *s^H^*, where *H* is the Hurst exponent, irrespective of the order of the detrending polynomial. In the simple case of fully random (”white noise”) increments *H* = 1*/*2, while *H >* 1*/*2 correspond to positively and *H <* 1*/*2 to negatively correlated increments, respectively.

While DFA is known for its robustness on long time scales, providing reliable estimates up to *s* ≲ *L/*4, it is not applicable on short time scales, due to exact or near-exact fitting problem, and even for noise-free simulated data approaches the asymptotic behavior only at *s* ≳ 8. The latter is the reason why DFA is often combined with WTA to cover all scales (for deeper details see, e.g. [41, 42] and references therein). In the WTA procedure, the observational data series *x_i_* is being splitted into windows of size *s*, and local sums 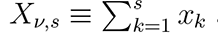 are calculated for each window *ν*, resembling the standard FA procedure. In high-order WTA*p*, further detrending is based on the analysis of the statistics of the *p*-th differences of the local sums *X_ν,s_*. The latter results in WTA being considerably less noise-robust than DFA, especially at large scales, providing relevant estimates only at 1≤*s* ≲ *L/*100, according to the results of simulation-based studies (see, e.g., [41] and references therein).

In their conventional implementations, both DFA and WTA procedures consider non-overlapping time windows resulting in discontinuity in the series of residuals, a common drawback that has been reduced in the central moving average (CMA) based detrended fluctuation analysis [24], where local polynomial fits *p_ν,i_*have been replaced by the local averages 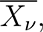 and non-overlapping windows of length *s* by moving windows of the same size, such that the residuals are calculated as 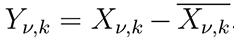. In terms of scale range and noise robustness, CMA occupies an intermediate position between DFA and WTA methods, and thus could be considered as an all-scale substitute to the combination of the above two methods. In surrogate data, CMA provides with reasonable approximations to asymptotic behavior already at small scales at least for odd window sizes, such that there is no bias of the central point in the window due to discreteness.

Since individual dendrochronological series are characterized by different lengths and locations on the time scale representing life cycles of each tree, and thus only partially overlap with each other, to avoid further loss of data from splitting into non-overlapping windows, in our study, we applied both DFA and CMA is gliding windows with single-year steps.

#### 2.5.2. Detrended Cross-Correlation Analysis

To quantify the relative dynamics, the above DFA (or CMA) procedures can be generalized to the analysis of several simultaneously observed time series 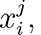 pairwise covariances between the residuals series (after subtraction of the average or polynomial fit, respectively) being calculated as [22, 23]

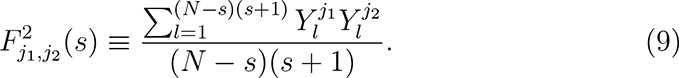

Next, for all *j*_1_*, j*_2_ = 1*, . . ., m* one obtains the covariance matrix

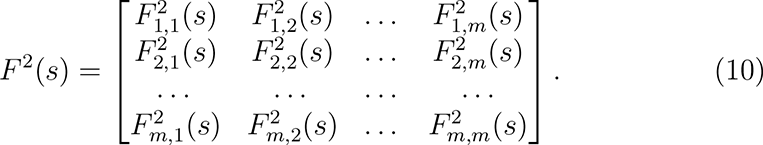

While the diagonal elements of the matrix 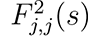 are simple variances and correspond to the fluctuation functions *F* ^2^(*s*) in the conventional detrended fluctuation analysis (DFA) [14], non-diagonal elements of the fluctuation matrix 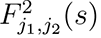 represent cross-covariances. To obtain cross-correlations between data series *j*_1_ and *j*_2_ (for example, between the annual tree-ring data and hydrometeorological variations), one calculates [22, 23]

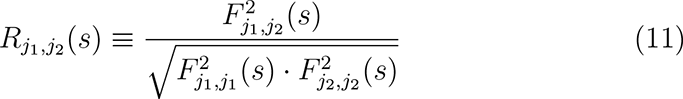

and obtain the matrix of cross-correlation coefficients by normalization

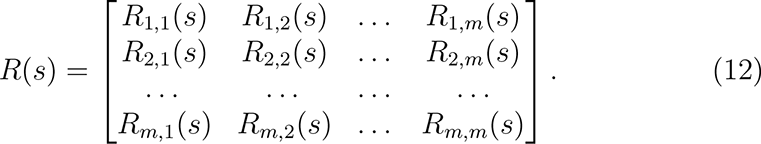

Finally, to exclude the spurious correlations induced by cross-modulation of the analyzed time series, one can also obtain partial correlation coefficients by calculating the inverse of the cross-correlation coefficient matrix

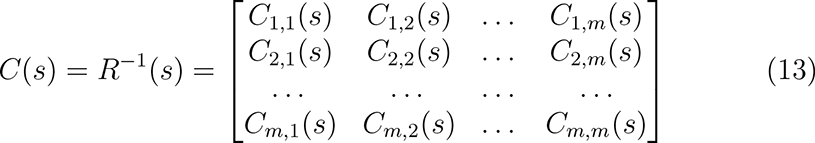

followed by its normalization as

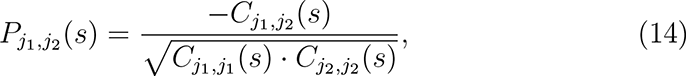

where the latter coefficients characterize intrinsic correlations between data series *j*_1_ and *j*_2_.

In the following, we apply a similar procedure to the fluctuations obtained by the CMA method.

Finally, to account for possible delayed response of the trees growth rates to hydroclimatic variations, we analyze not only synchronous, but also delayed cross-correlations [43] with time lages up to five years, corroborating the hypothesis that annual tree-ring width could be associated with any of the monthly climate variations over the preceeding 60 months, from September to August (since tree growth in the considered climate zone typically occurs only between May and August).

Figure 2 exemplifies the CMA-based multi-scale detrended cross-correlation analysis procedure. In this example, domination of negative correlations between the PDSI variations taken for August each year and a single TRW data series for tree Nr. 115, at scale *s* = 5, with 2-year delay lag is demonstrated. The covariances *F_ij_* and cross-correlation coefficients *R_ij_* are next calculated by averaging over *all* positions of the local time window of the same duration *s* = 5 years and 2-year time lag between the analyzed records.

**Figure 2:**
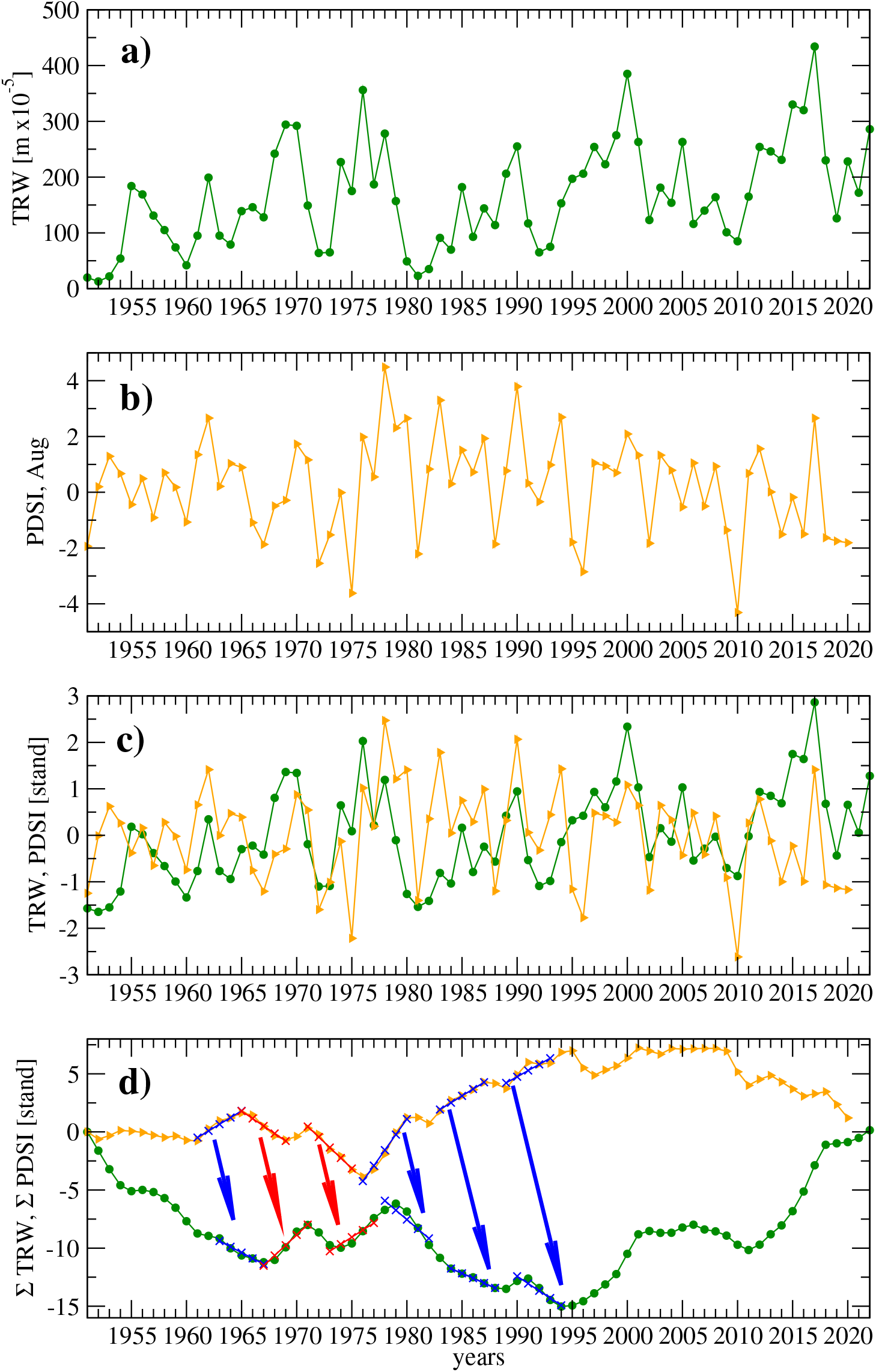
Simple illustration of the CMA-based multi-scale detrended cross-correlation analysis procedure. (a) TRW and (b) PDSI data series are first standardized by subtracting their averages and dividing by standard deviations (c). Next, cumulative profiles (d) are calculated, and local windows of size *s* are determined. In each window, the local mean is calculated and subtracted, resulting in the estimate of the relative trend around the local mean. Panel (d) exemplifies the correspondence between the variations in the PDSI index with local increasing (blue) and negative (red) trends indicating shifts towards more humid and more dry conditions, respectively, and local trends in a single representative TRW profile corresponding to either increasing (red) or decreasing (blue) annual growth rate with two-year delay.

### 2.6. Statistical Significance Testing

To reveal statistically significant discrepancies between measurements obtained for different local subareas, we employed the one-way non-parametric ANOVA (Kruskal-Wallis) statistical test with significance threshold at *p* = 0.05. In multi-group comparison scenarios, in order to specify particular contributors to the overall discrepancies, we also performed post-hoc analysis using Mann-Whitney U-test with the same confidence threshold level.

## 3. Results

### 3.1. Multispectal Remote Sensing

We used multispectral imaging to characterize the local hydrological conditions that in turn also affect the overall vegetation. Figures A.13 and A.14 in the Appendix A show that the vegetation indices (EVI, RTVICore, MTVI2) generally follow the elevation profile, with highest values observed for the elevated dry land area surrounding the peat bog, and reduced values for the peat bog, with a tendency of monotonous reduction with lowering elevation both within and outside of the peat bog. The NDWI index (also represented in Fig. 1), as expected, shows the reciprocal pattern, with a clear trend indicating higher water content with lowering. Figure A.14 shows boxplots indicating discrepancies between the local measurements in each of the four studied areas (western, middle and eastern parts of the peat bog, with continuously reduced elevation, as well as dry land areas at surrounding elevations), including five multispectral bands (Blue, Green, Red, RedEdge, NIR), as well as calculated vegetation indices (EVI, NDWI, RTVICore, MTVI2). Statistically significant discrepancies are indicated by (red) annotation, with reported *p*-values according to one-way non-parametric ANOVA (Kruskal-Wallis) statistical test, while particular locations where respective measurements indicated significant discrepancies with the dry land (according to the post-hoc test results) are denoted by (red) asterisks.

### 3.2. Tree-Ring Data and Hydroclimatic Time Series

Figure 3 shows dendrochronological data series (averages over four groups corresponding to three locations within the peat bog area and the surrounding dry land area, with the latter acting as relevant control, are provided), as well as historical temperature and PDSI data. While the full duration of chronologies are indicated in the stacked barplot in Fig. 4, Fig. 3 shows data series only for the time frame since 1901, when significant number of tree chronologies in each local area, as well as climate variations data are available. The figure shows that the dry land TRW data exhibits pronounced non-stationarity, including two major bursts associated with the renewal of considerable fractions of trees. We hypothesize that the replacement of the older trees by the new ones could be enforced by the loss of a significant fraction of trees during two major climate extremes, a prolonged drought in the early 1930s characterized by near 10-year decline in PDSI with culmination in 1936 (accompanied by several hot weather anomalies), and a major winter temperature anomaly in 1941-1942 (with night temperatures registered at the local meteo station going below 45*^o^*C in late January) [44], the latter resulting in 3-10-fold growth suppression in the majority of the survived trees in the same year. Remarkably, no similarly pronounced effect could be observed for the trees located within the peat bog area. For a more relevant comparison, separate curves for the TRW data for chronologies starting before 1930 and 1940, respectively, indicate that in the following years they exhibit similar variability patterns like the overall average in this dry area, and thus likely demonstrate similar cross-correlation patterns with meteorological variatibility data (shown in the middle and lower panels of Fig. 3).

**Figure 3:**
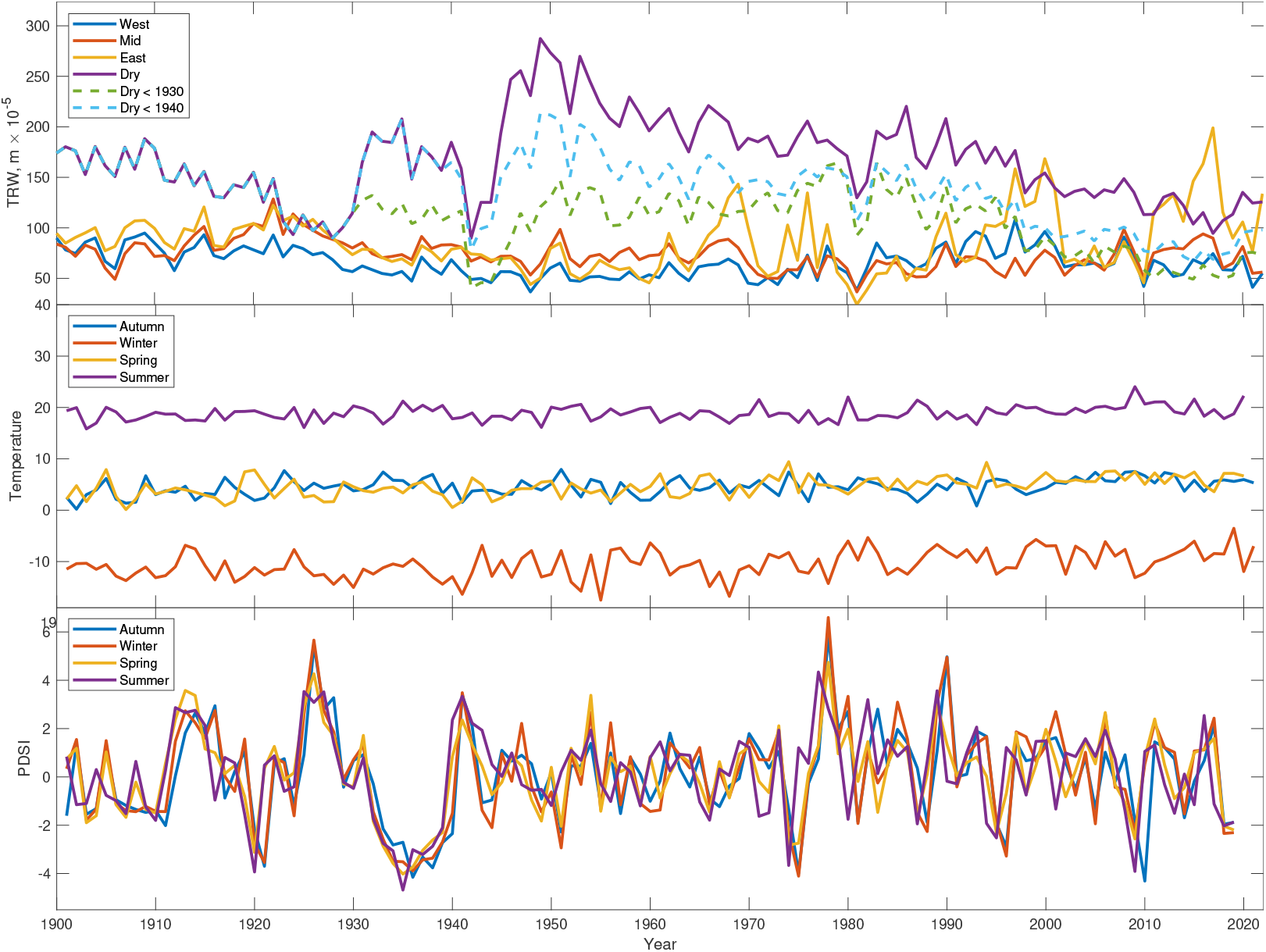
TRW data averaged over four different subareas: three locations in the peat bog area, altogether 101 data series, as well as averaged data over 40 data series from three dry land locations at local elevations surrounding the peat bog area (upper panel). Seasonal average temperature (middle panel) and PDSI (lower panel) variations. Since in the dry area a large fraction of trees emerged between 1930s and 1950s, dashed curves additionally show data for chronologies starting *<* 1930 and *<* 1940, respectively.

**Figure 4:**
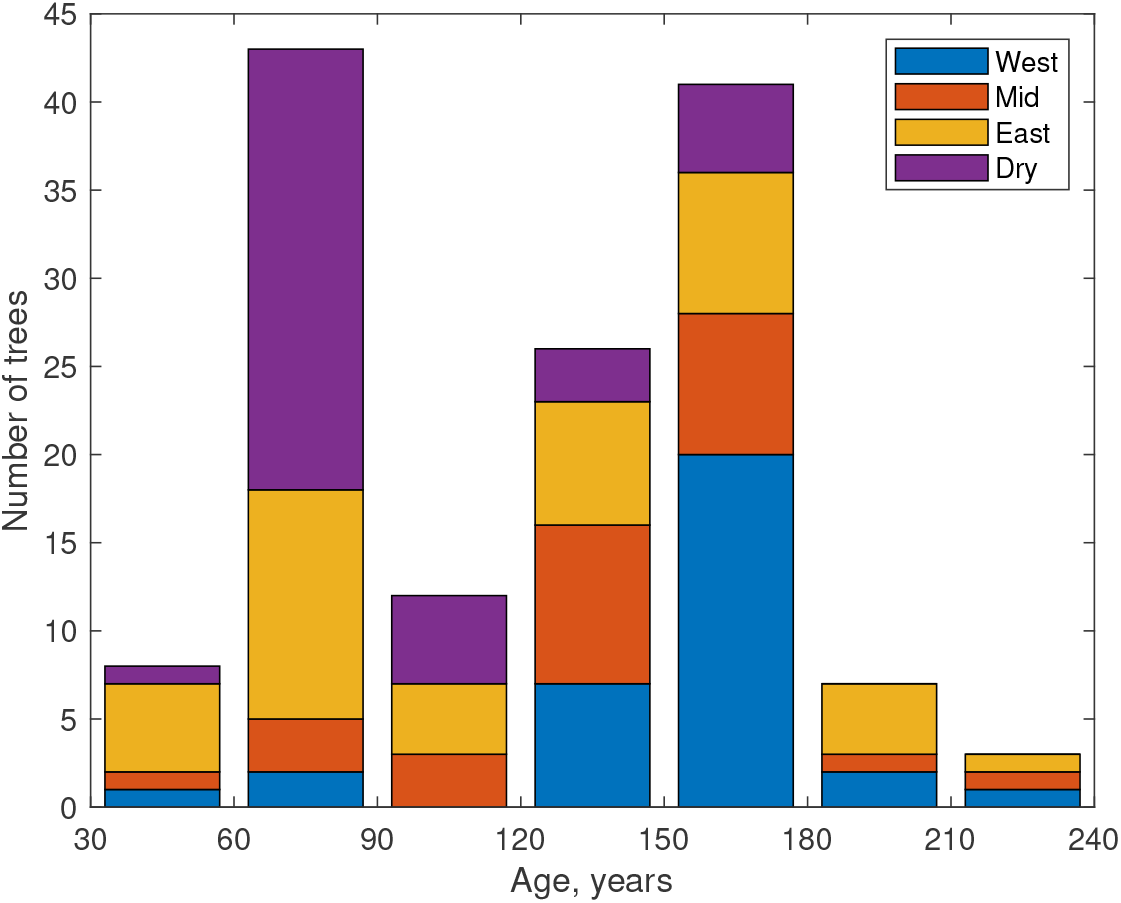
Stacked barplot representing the distribution of the durations of dendrochronologies for each of the studied locations.

### 3.3. Detrended Fluctuation Analysis

Figure 5 shows fluctuation functions for the TRW, temperature and PDSI data variations, obtained by first- and second-order DFA methods, as well as by CMA detrended fluctuation analysis methods. The figure shows that for the TRW data *all* three methods provide estimates *H* ≅ 1.0 ± 0.1, that is also in a reasonable agreement (taking into account the length and temporal resolution of the records) with results for tree-ring data reported in recent literature [19, 20, 42]. For the temperature records, all methods show asymptotic slopes *H* between 0.6 and 0.7, that is also in agreement with earlier results for hydroclimatic time series (*H* ≅ 0.65 appeared the most commonly observed exponent for long-term temperature observations) [15, 45, 41]. Regarding the PDSI, there is a characteristic crossover, indicating additional short-term memory corresponding to the accumulation of the drought effects, although the asymptotic slopes are likely in the similar range as for the temperature data, that appears in between the typical exponents for the (non-accumulated) precipitaion and (accumulated) river runoff data [46, 47] (for a recent review, see also [48] and references therein).

**Figure 5:**
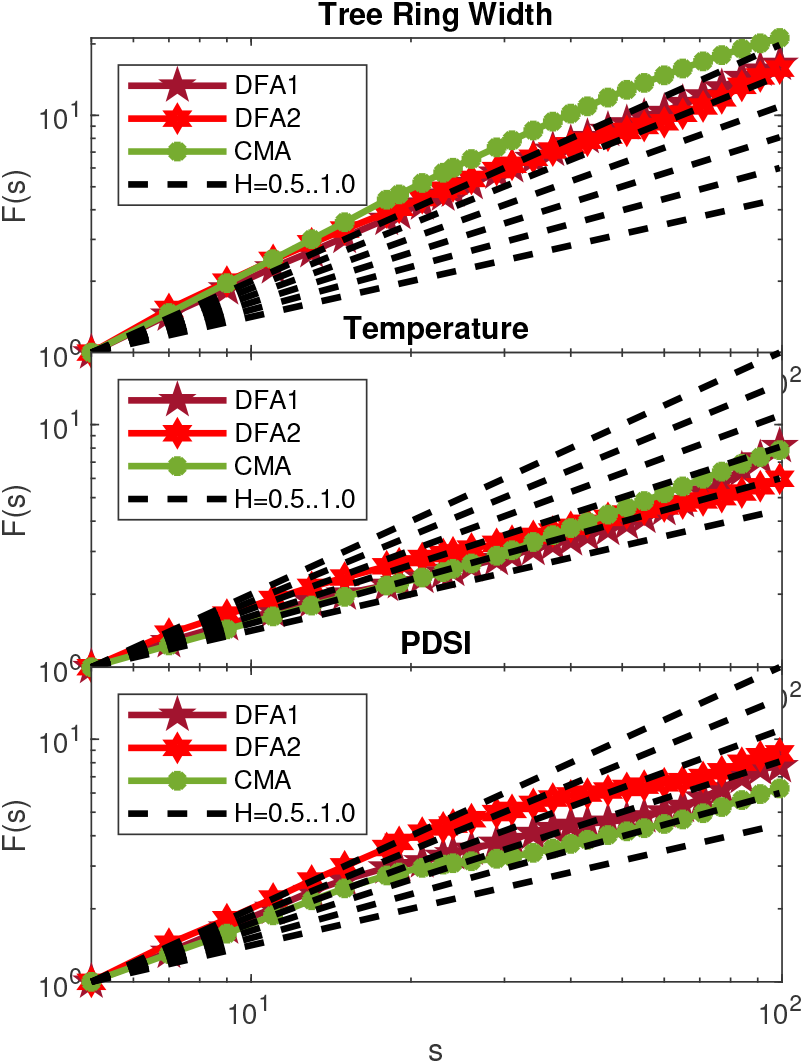
Fluctuation functions for the TRW, temperature and PDSI data variations, obtained by first- and second-order DFA methods, as well as by CMA detrended fluctuation analysis methods. Dashed lines show theoretical slopes corresponding to linearly long-term correlated (fractional Gaussian noise) data series *F* (*s*) ∝ *s^H^*, from the simplest case *H* = 0.5 (corresponding to uncorrelated “while noise” data) to *H* = 1.0 (corresponding to the upper bound for stationary data).

### 3.4. Detrended Cross-Correlation Analysis

In the following, we analyze cross-correlations between individual dendrochronological data series, using both conventional cross-correlation coefficient matrix and the multi-scale cross-correlation coefficients obtained by DPCCA with CMA- and DFA-based detrending calculated for scales *s* = 2*^k^* + 1, where *k* = 1, 2*, . . .,* 5. Figure 6 indicates that at all scales highest positive cross-correlations could be observed for the trees located within the areas with minumum and maximum elevations, in the lowest area in the eastern part of the peat bog, and in the elevated dry land area west of the peat bog, while only weak negative correlation tendency could be observed between these areas.

**Figure 6:**
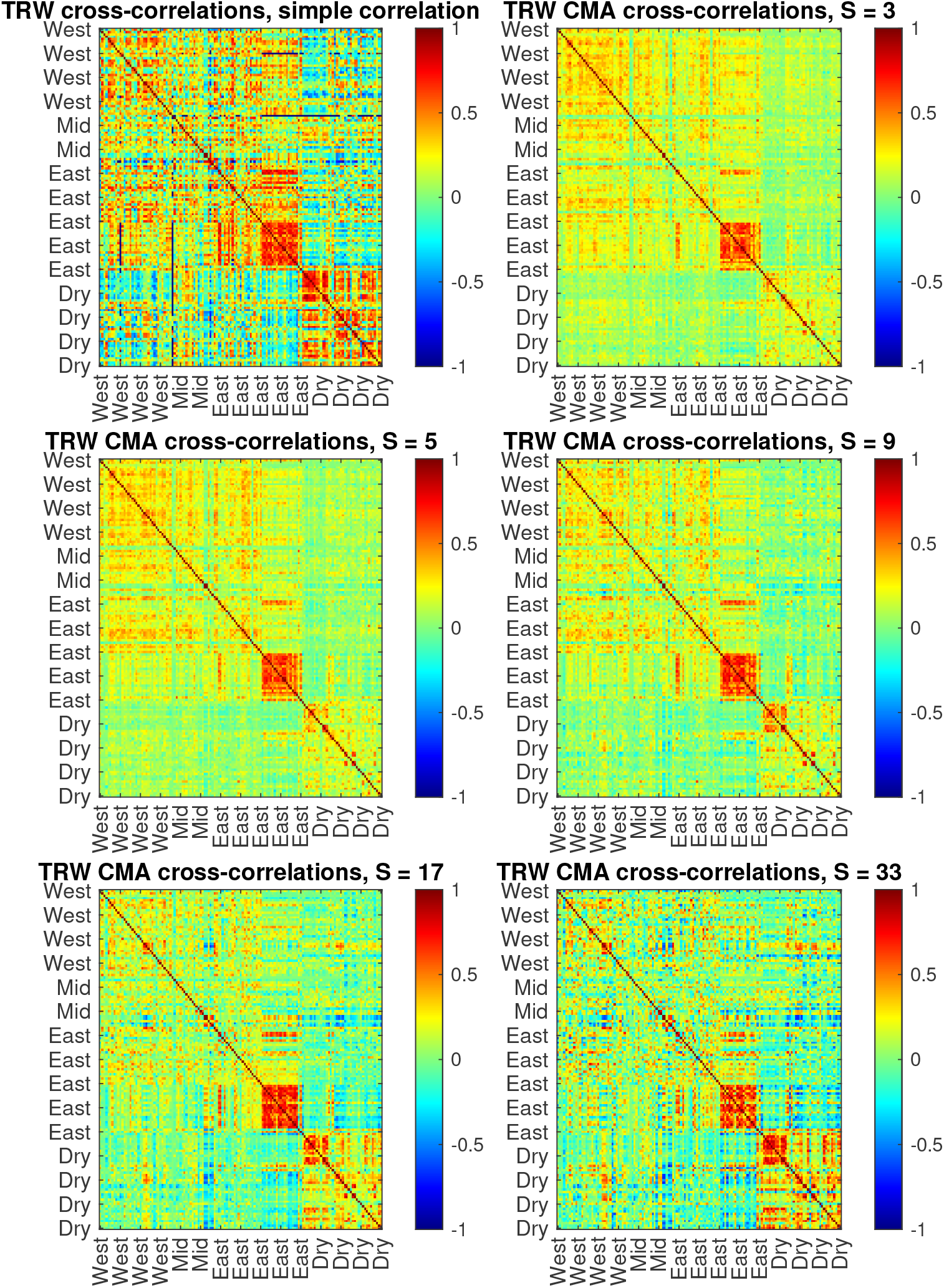
Multi-scale cross-correlations between individual TRW data series, altogether for *n* = 140 trees growing in different parts of the bog (*n* = 32 in the western, *n* = 26 in the central, *n* = 42 in the eastern), as well as in the surrounding elevated dry land area (*n* = 40 altogether in three surrounding locations). The upper left panel shows the conventional cross-correlation coefficient matrix, while the remaining panels show the multi-scale DPCCA cross-correlations (with CMA-based detrending) calculated for scales *s* = 2*^k^* + 1, where *k* = 1, 2*, . . .,* 5.

Next, we analyze cross-correlations between individual dendrochronological data series and the local temperature variations within the 5-year long time window up to the end of the growth period for the year of interest (from September to August). Although we have initially analyzed monthly temperature variations, in order to reduce the impact of single outliers in the correlation patterns (that are barely significant when considering appropriate corrections for multiple testing), we show the heatmaps after 3 × 3 moving average, thus reducing noise in the images.

Figures 7 and 8 indicate that TRW are negatively correlated with spring and summer temperatures and positively correlated with the Palmer drought severety index (PDSI) in the same year, indicating that heatwaves and droughts represent the limiting factors for the trees growth. However, regarding more long-term effects, remarkable contrasts can be observed between areas with different local hydrological conditions already at interannual scales. In particular, for the sphagnum bog area positive TRW trends over several consecutive years tend to follow negative PDSI trends and positive spring and summer temperature trends of the same duration with a time lag typically between one and three years, indicating that prolonged dry periods, as well as warmer springs and summers appear beneficial for the increased annual growth. In contrast, for the surrounding elevated dry land area nearly reversed tendency can be observed, with pronounced negative long-term correlations with temperature and positive correlations with PDSI.

**Figure 7:**
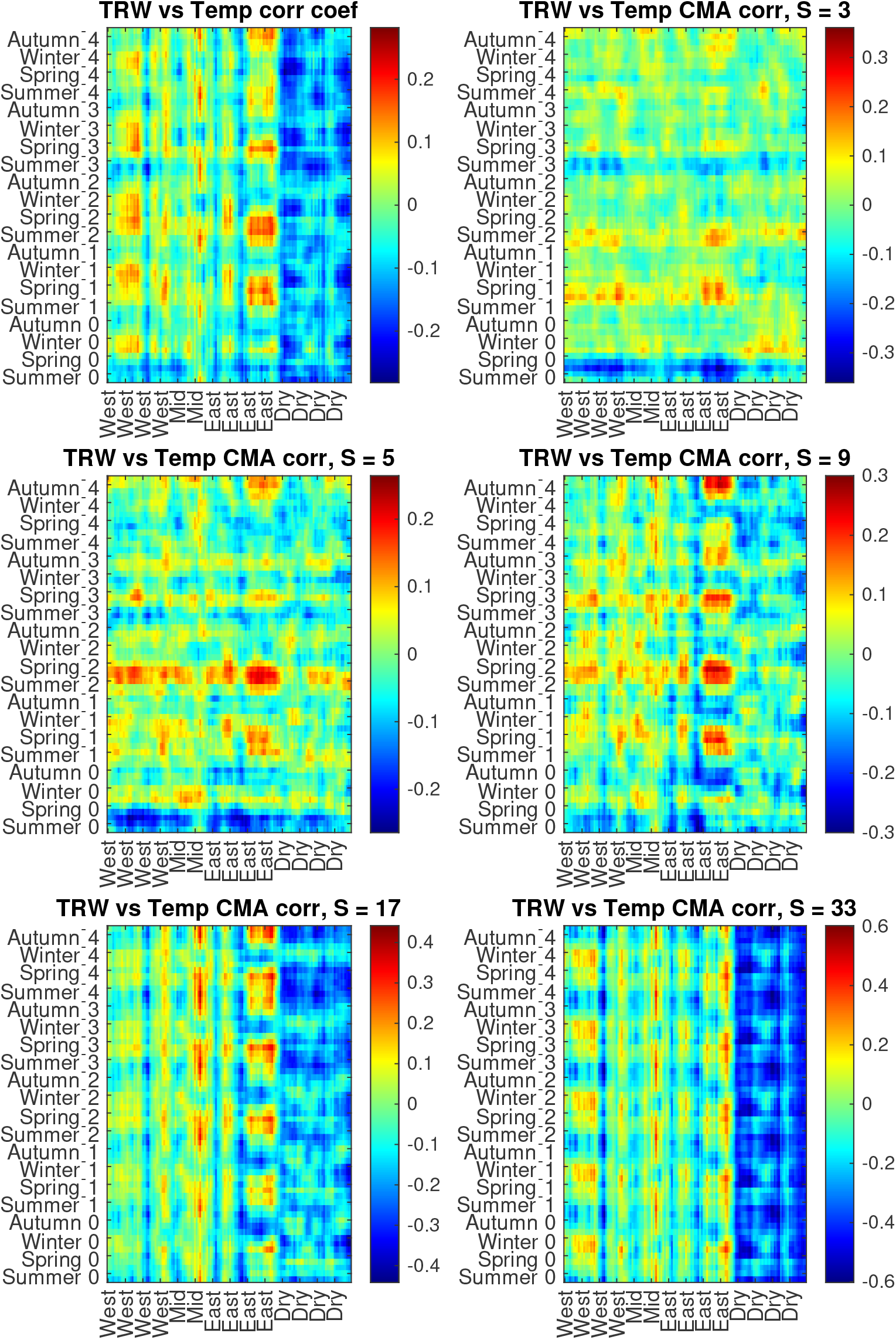
Multi-scale conventional and DPCCA (with CMA-based detrending) cross-correlation coefficients between individual TRW data series, altogether for *n* = 140 trees growing in different parts of the bog (*n* = 32 in the western, *n* = 26 in the central, *n* = 42 in the eastern), as well as in the surrounding elevated dry land area (*n* = 40 altogether in three surrounding locations), and the local temperature variations over five years prior and during the growth period for a given year (September to August). Monthly resolution data were analyzed, but for noise reduction and better visibility displayed with seasonal resolution obtained by moving average in a 3 × 3 window over the entire heatmap.

**Figure 8:**
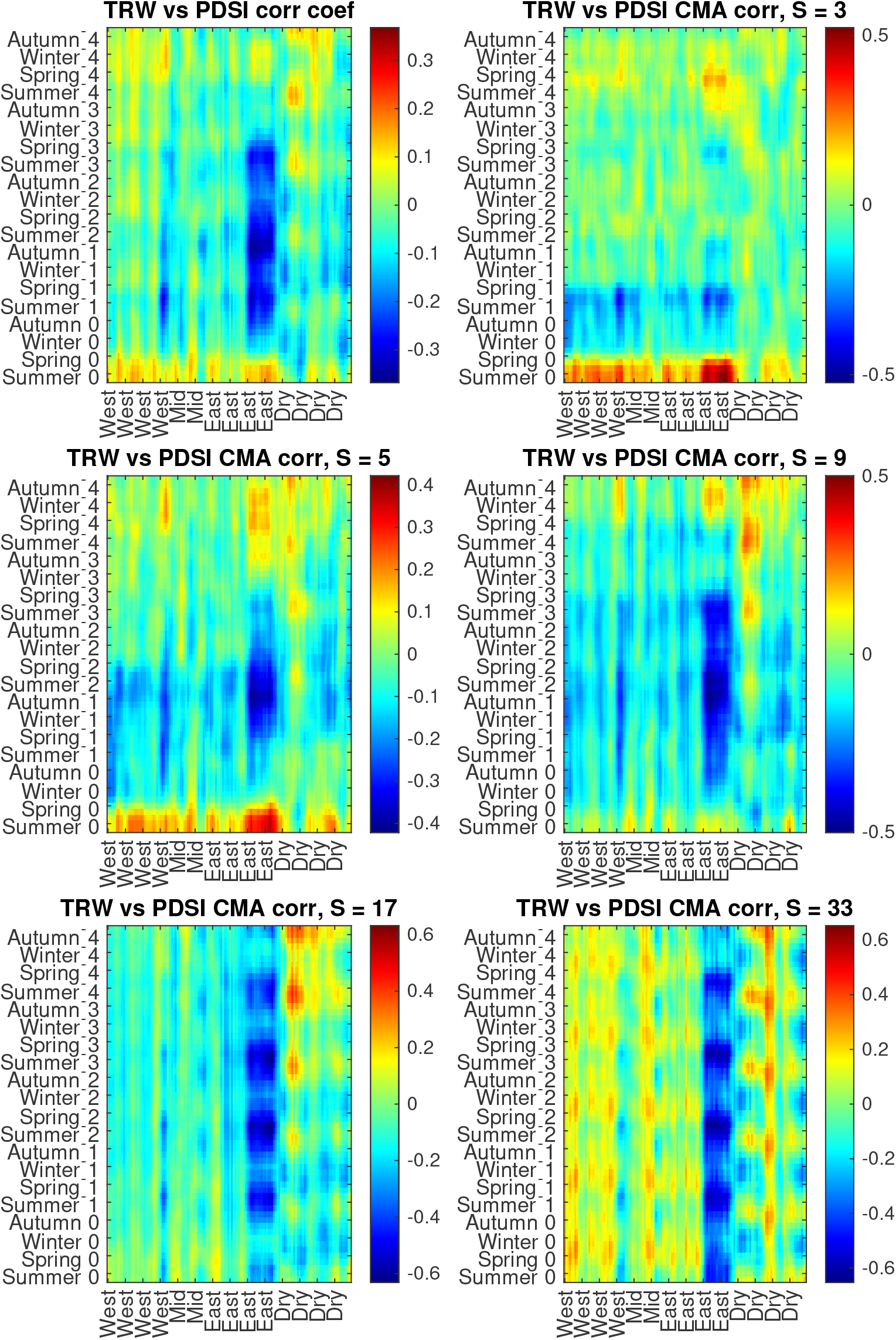
Multi-scale conventional and DPCCA (with CMA-based detrending) cross-correlation coefficients between individual TRW data series, altogether for *n* = 140 trees growing in different parts of the bog (*n* = 32 in the western, *n* = 26 in the central, *n* = 42 in the eastern), as well as in the surrounding elevated dry land area (*n* = 40 altogether in three surrounding locations), and the local PDSI variations over five years prior and during the growth period for a given year (September to August). Monthly resolution data were analyzed, but for noise reduction and better visibility displayed with seasonal resolution obtained by moving average in a 3 × 3 window over the entire heatmap.

However, one can easily notice that long-term negative correlations between TRW and temperature variations for the elevated dry land area that only become stronger with increasing scale, and thus could be potentially attributed to the long-term increase of temperatures associated with climate change, also eventually coincide with the overall reduction of the trees growth rate in this area, that is reminiscent to a similar pattern around 100 years ago, and thus could be also at least in some part attributed to the ageing effects of the trees (see Fig. 3). Since in all cases the respective trends could be in the first approximation represented by their linear fits, we next repeat similar analysis using first-order DFA for detrending. Figure 9 indicates that in the modified DPCCA results these previously pronounced negative correlations have been considerably reduced, and remain notable for the spring temperatures only. Figure 10 also shows that, while the absolute values of cross-correlation coefficients have been reduced after linear detrending, local contrast patterns between correlations in the lowest area in the eastern part of the peat bog area and its surroundings have been only enhanced.

**Figure 9:**
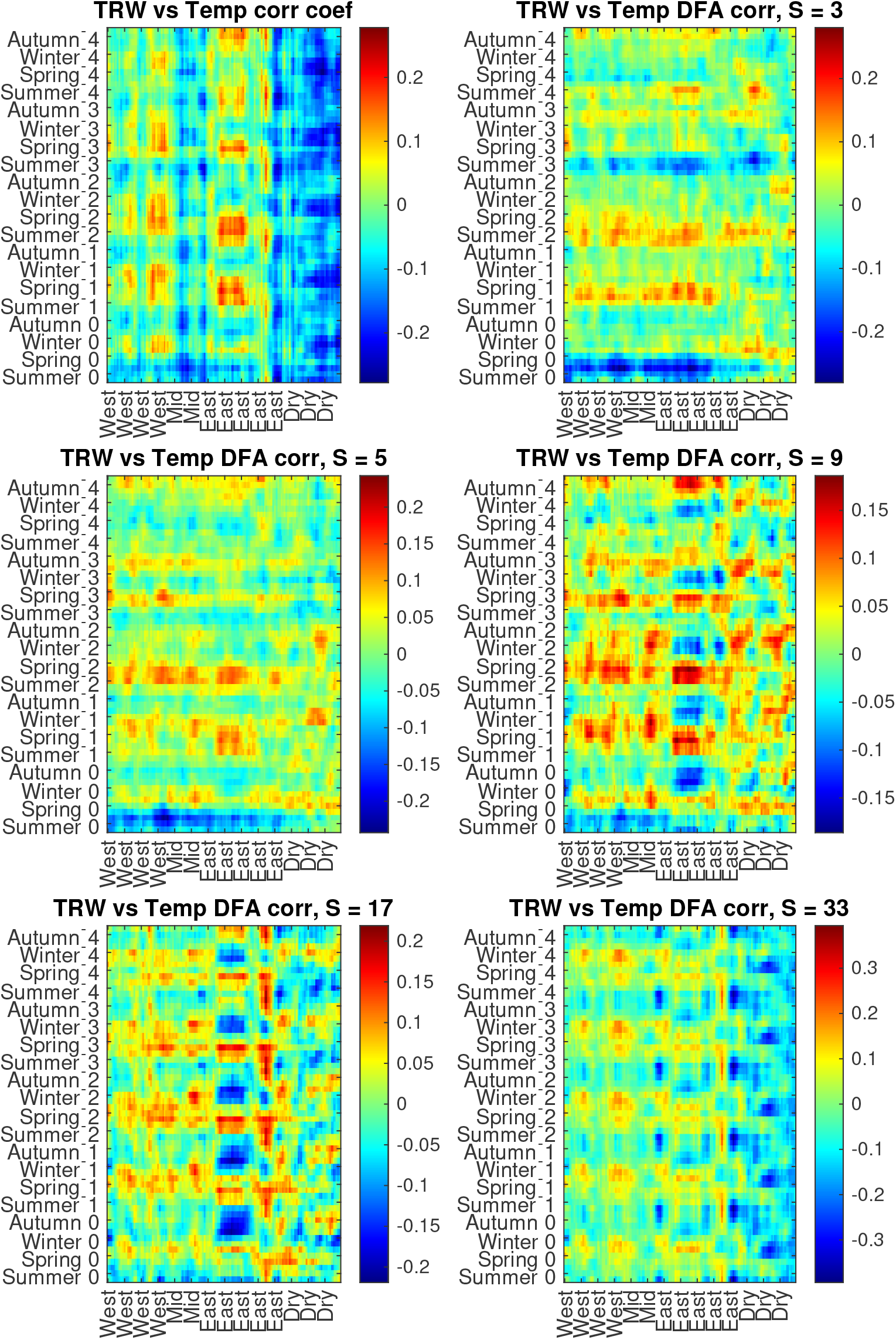
Multi-scale conventional and DPCCA (with DFA-based linear detrending) cross-correlation coefficients between individual TRW data series, altogether for *n* = 140 trees growing in different parts of the bog (*n* = 32 in the western, *n* = 26 in the central, *n* = 42 in the eastern), as well as in the surrounding elevated dry land area (*n* = 40 altogether in three surrounding locations), and the local temperature variations over five years prior and during the growth period for a given year (September to August). Monthly resolution data were analyzed, but for noise reduction and better visibility displayed with seasonal resolution obtained by moving average in a 3 × 3 window over the entire heatmap.

**Figure 10:**
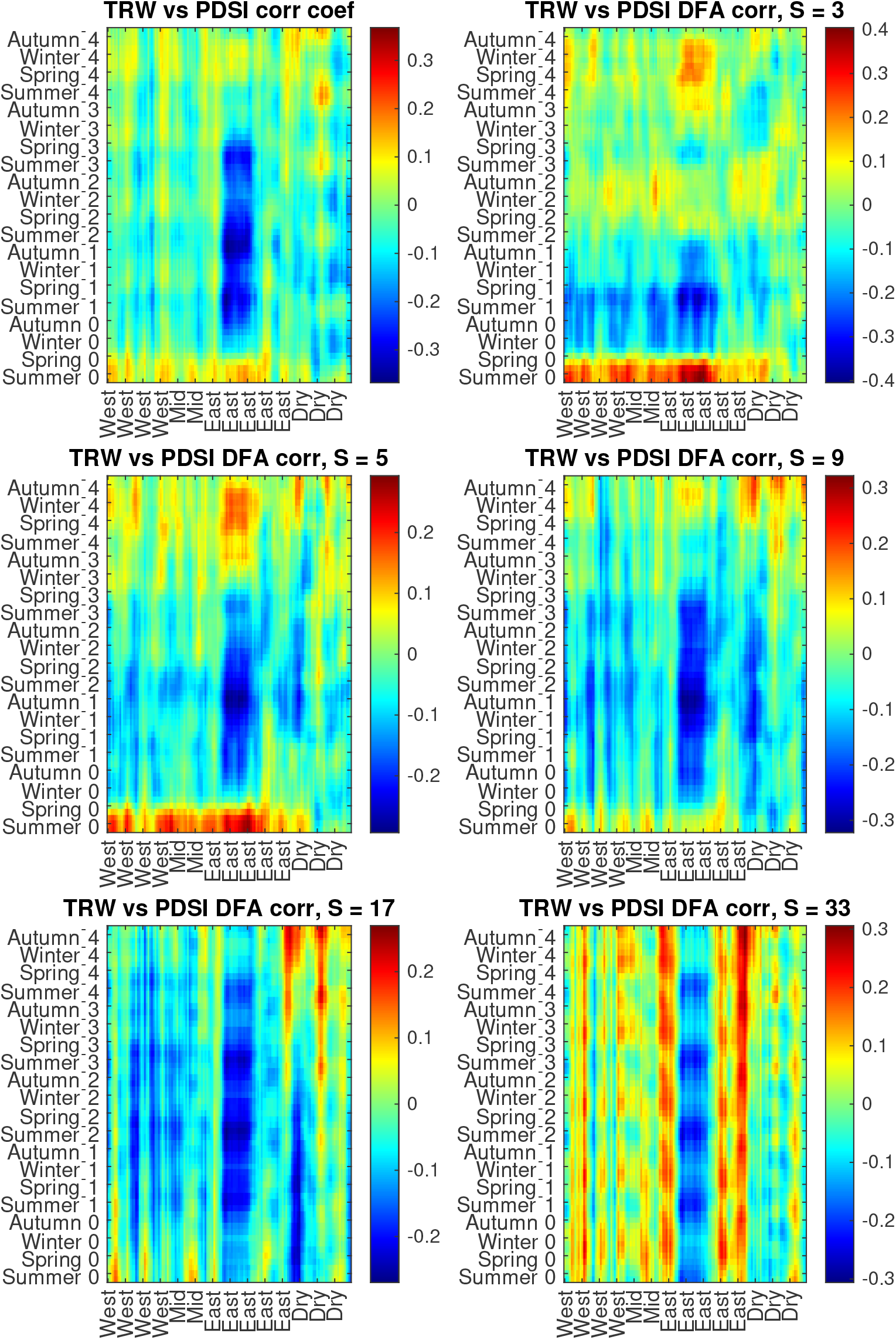
Multi-scale conventional and DPCCA (with DFA-based linear detrending) cross-correlation coefficients between individual TRW data series, altogether for *n* = 140 trees growing in different parts of the bog (*n* = 32 in the western, *n* = 26 in the central, *n* = 42 in the eastern), as well as in the surrounding elevated dry land area (*n* = 40 altogether in three surrounding locations), and the local PDSI variations over five years prior and during the growth period for a given year (September to August). Monthly resolution data were analyzed, but for noise reduction and better visibility displayed with seasonal resolution obtained by moving average in a 3×3 window over the entire heatmap.

To further validate the ageing hypothesis, we also consider partial correlations using modified DPCCA but with CMA based detrending which does not eliminate linear trends from the data, for the annual observations of TRW, tree age, and climate variations (temperature or PDSI, respectively), in order to eliminate linear associations with the tree age from cross-correlations with the climate variations, separately for each of the studied trees. Figures 11 and 12 indicate that the effects are qualitatively similar to those when replacing CMA by linear detrending DFA, while the keynote effects reported above become only further highlighted by higher absolute values of cross-correlation coefficients, indicating the major impact of the local hydrological conditions on the drought stress resilience.

**Figure 11:**
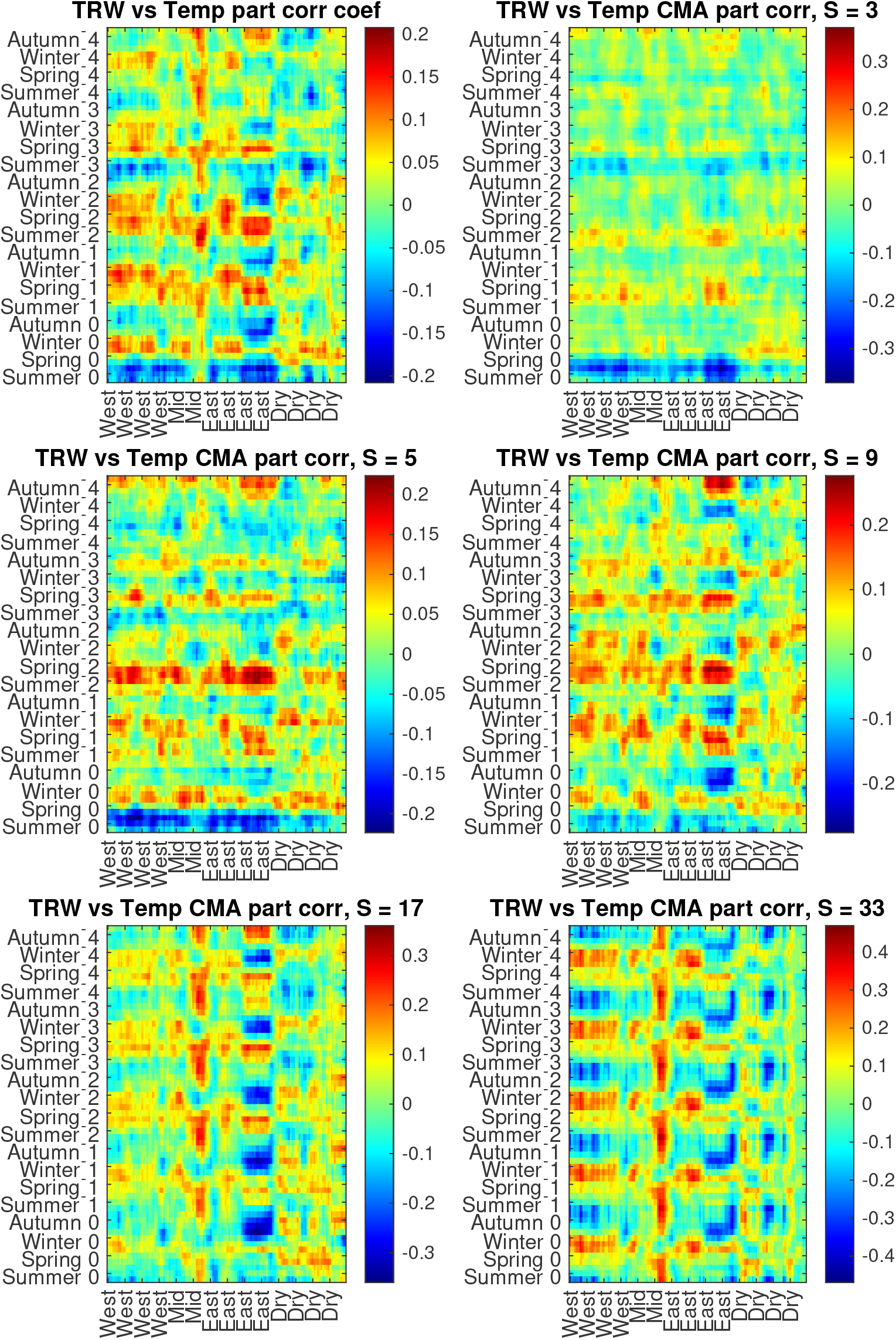
Multi-scale conventional and DPCCA (with CMA-based detrending) partial cross-correlation coefficients between individual TRW data series, altogether for *n* = 141 trees growing in different parts of the bog (*n* = 33 in the western, *n* = 26 in the central, *n* = 42 in the eastern), as well as in the surrounding elevated dry land area (*n* = 40 altogether in three surrounding locations), and the local temperature variations over five years prior and during the growth period for a given year (September to August). Monthly resolution data were analyzed, but for noise reduction and better visibility displayed with seasonal resolution obtained by moving average in a 3×3 window over the entire heatmap.

**Figure 12:**
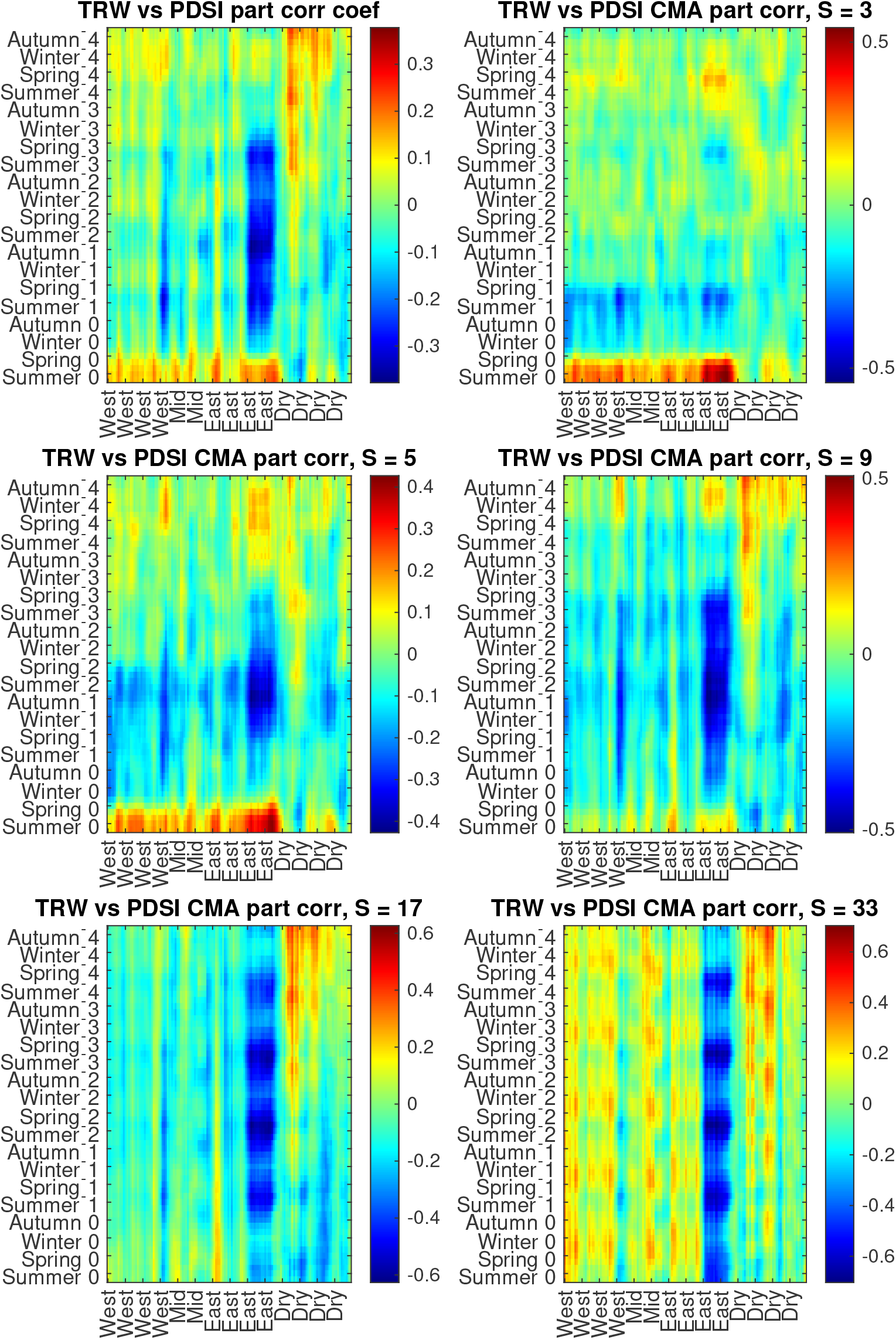
Multi-scale conventional and DPCCA (with CMA-based detrending) partial cross-correlation coefficients between individual TRW data series, altogether for *n* = 141 trees growing in different parts of the bog (*n* = 33 in the western, *n* = 26 in the central, *n* = 42 in the eastern), as well as in the surrounding elevated dry land area (*n* = 40 altogether in three surrounding locations), and the local PDSI variations over five years prior and during the growth period for a given year (September to August). Monthly resolution data were analyzed, but for noise reduction and better visibility displayed with seasonal resolution obtained by moving average in a 3×3 window over the entire heatmap.

## 4. Discussion, Conclusion, and Outlook

Recent and ongoing climate alterations are associated with changes in the forest structure and composition that appear especially pronounced in the northern latitudes, strongly affecting the southern edge of the boreal ecotone [49]. In recent decades, exacerbating heat- and drought stress conditions already resulted in significant alternations in the boreal forest ecosystems both in Eurasia and North America, including changes in demographic rates, disturbance regimes, and range shifts of some tree species [50, 51]. There are certain indications that in some areas the southern edge of the boreal ecotone already exhibits a northwards shift [52], and there are raising concerns that these alterations could act as a feedforward driver reinforcing the effects of global climate warming at least on the local and regional scales [53].

Despite recent evidence that *Pinus* are among the few species in the boreal ecosystem responding favorably to increasing temperatures [49], they are particularly sensitive to prolonged droughts, and thus temperature effects likely play a secondary role, with warming promoting better growth only during non-drought years [54, 55]. Therefore, local hydrological conditions play a major role in the stress resilience and thus also in the preservation of the boreal ecosystems, along with global and regional climate drivers, not only directly but also via complex interspecies interactions in the ecosystem [56, 57]. Accordingly, a better understanding of the complex interplay of multiple factors affecting stress resilience of vulnerable forest ecosystems is essential for their efficient management, that in turn requires application of relevant measurement techniques and adequate mathematical methods.

While long-term climatological, dendrological, hydrological, meteorological, and remote sensing based measurements provide extensive information on the state of the ecosystem and its potential drivers, natural observations are typically characterized by long-term persistence [8, 58, 9, 59], considerably affecting the interpretation of the statistics provided by seemingly established mathematical methods [15, 60, 46, 47, 41]. In some cases, this has already led to spurious results, such as the underestimation of clustering effects in tree-ring based climate reconstructions compared to available long-term series of direct measurements [10], as well as overestimation of the significance of local trends in climate variations [11, 12].

To overcome the above limitations, in recent years detrended fluctuation analysis methods, especially DFA and WTA have been recommended as appropriate tools to analyze long-term tree-ring data and respective climate reconstructions, since they are capable of overcoming inevitable drawbacks of conventional correlation analysis when applied to long-term correlated data series, including non-stationary regimes [19, 20, 42]. In this work, we follow this direction and apply the recently proposed DPCCA method [23] to analyze cross-correlations between tree-ring data and climate variations.

While in earlier dendrochronological studies [19, 20, 42] second-order detrending has been commonly applied, we believe that in the context of the associations between the TRW data and interannual climate variations cross-correlations between residuals after second-order polynomial detrending is a quantity that is quite difficult to interpret. Furthermore, fluctuation analysis revealed that for the annual TRW data, as well as for the interannual temperature and PDSI variations (that are, on the one hand, already free of seasonal trends, while on the other hand, relatively short in terms of the number of samples available) CMA-based detrending and first-order DFA (being applied in a gliding window, in a similar way as CMA) provide with close estimates of correlation exponents as the second-order DFA (see Fig. 5), thus appropriately accounting for the autocorrelation patterns in the respective data series. In contrast, cross-correlations obtained by the modified DPCCA with CMA-based detrending and additional delayed response analysis could be easily illustrated in terms of relations between downward/upward interannual trends in the respective data series (see Fig. 2), and thus could be easily interpreted also by a domain expert in a non-mathematically focused field.

The studied forest ecosystem is located in a compact area within the Volzhsko-Kamsky State Nature Biosphere Reserve at the southern edge of the boreal ecotone that is particularly vulnerable to prolonged heat- and drought stress with a remarkable gradient in the local hydrological conditions, representing a prominent test bed for the investigation of the role of local hydrological factors in the climate stress resilience analysis. Our results indicate that, while the trees in the elevated dry land areas are generally characterized by higher growth rates, this comes at the cost of their considerably reduced drought stress resilience. Also more pronounced growth reduction with ageing, as well as more frequent replacement of the trees by the new ones, likely following significant losses of trees to climate extremes could be observed (see Fig. 3). In contrast, while exhibiting lower productivity, the forest ecosystem in the wetland area are characterized by longevity, with several chronologies dating back to 1780s - 1800s, several decades beyond the longest chronologies in the dry land area (see Fig. 4). They are also considerably less sensitive to heat- and especially drought stresses, and demonstrate considerably weaker ageing trends (see Fig. 3).

While tree-ring data are negatively correlated with spring and summer temperatures and positively correlated with PDSI in the same year indicating that heatwaves and droughts represent the limiting factors, at interannual scales remarkable contrasts can be observed between areas with different local hydrological conditions. In particular, for the sphagnum bog area positive TRW trends over several consecutive years tend to follow negative PDSI trends and positive spring and summer temperature trends of the same duration with a time lag between one and three years, indicating that prolonged dry periods, as well as warmer springs and summers appear beneficial for the increased annual growth. In contrast, for the surrounding elevated dry land area a reversed tendency can be observed, with pronounced negative long-term correlations with temperature and positive correlations with PDSI.

Finally, linear detrending and partial correlation analysis capable of distinguishing between ageing trends and climate stress response indicate that the observed associations with PDSI remain and in some cases are even further highlighted, indicating that drought stress represents the major challenge, while correlations with temperature are rather secondary, remaining significant only in certain seasons and play their role on shorter time scales.

We believe that the considered methodological modifications would be helpful in revealing the complex interconnections between global, regional and local factors to unravel and characterize in a quantitative manner the keynote factors governing stress resilience, and thus also welfare, vigor, and survival of the forest ecosystems, leading to improved risk assessment and thus also better environmental management planning.

## Acknowledgements

The authors would like to acknowledge the financial support of this research by the Russian Science Foundation (grant No. 22-76-10042), https://rscf.ru/project/22-76-10042/.

## Appendix A. Appendix

**Figure A.13:**
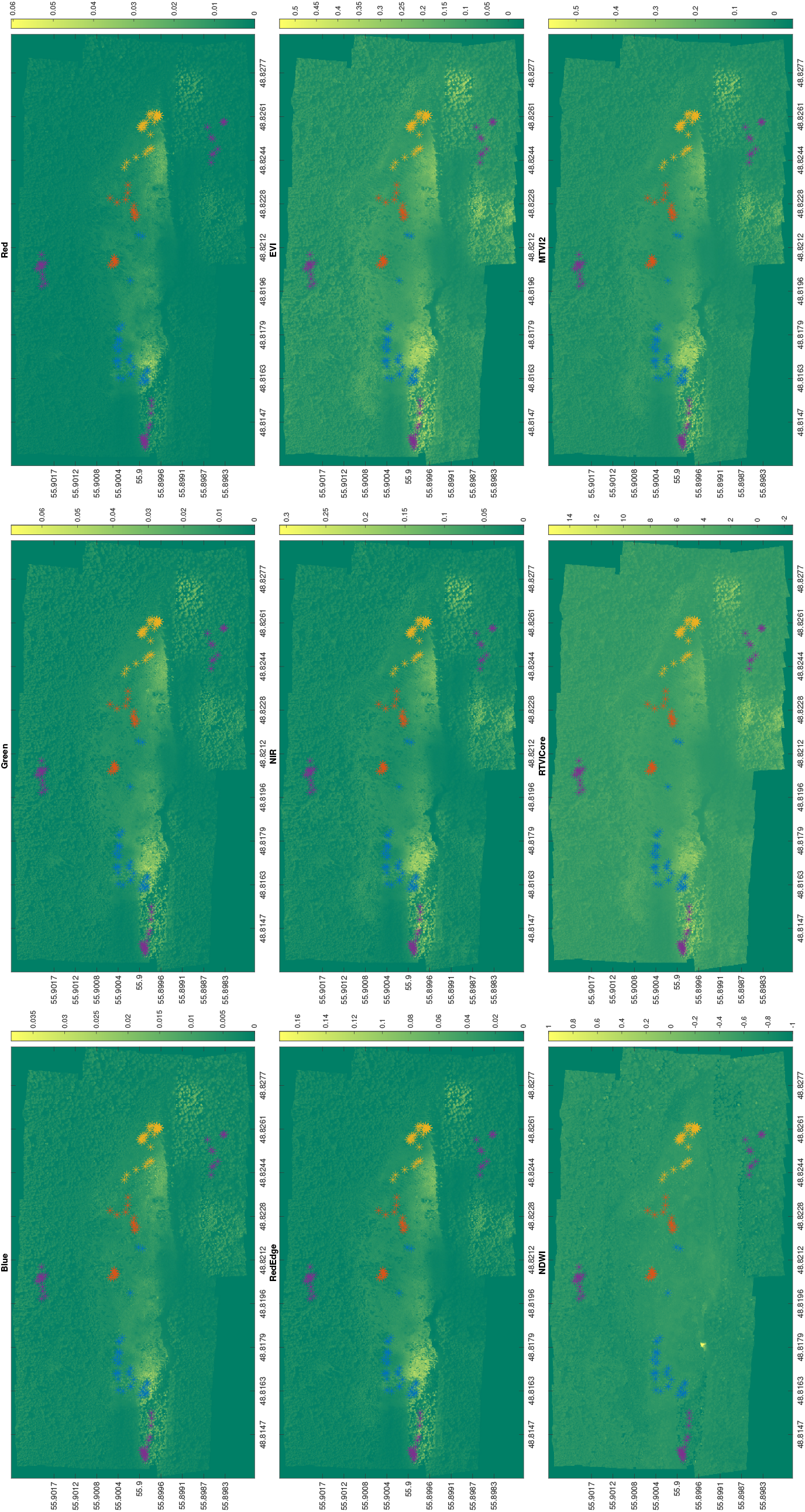
An overview of imaging data including five multispectral bands (Blue, Green, Red, RedEdge, NIR), as well as calculated vegetation indices (EVI, NDWI, RTVICore, MTVI2). Locations of the trees are denoted by colored asterisks (similar to Fig. 1).

**Figure A.14:**
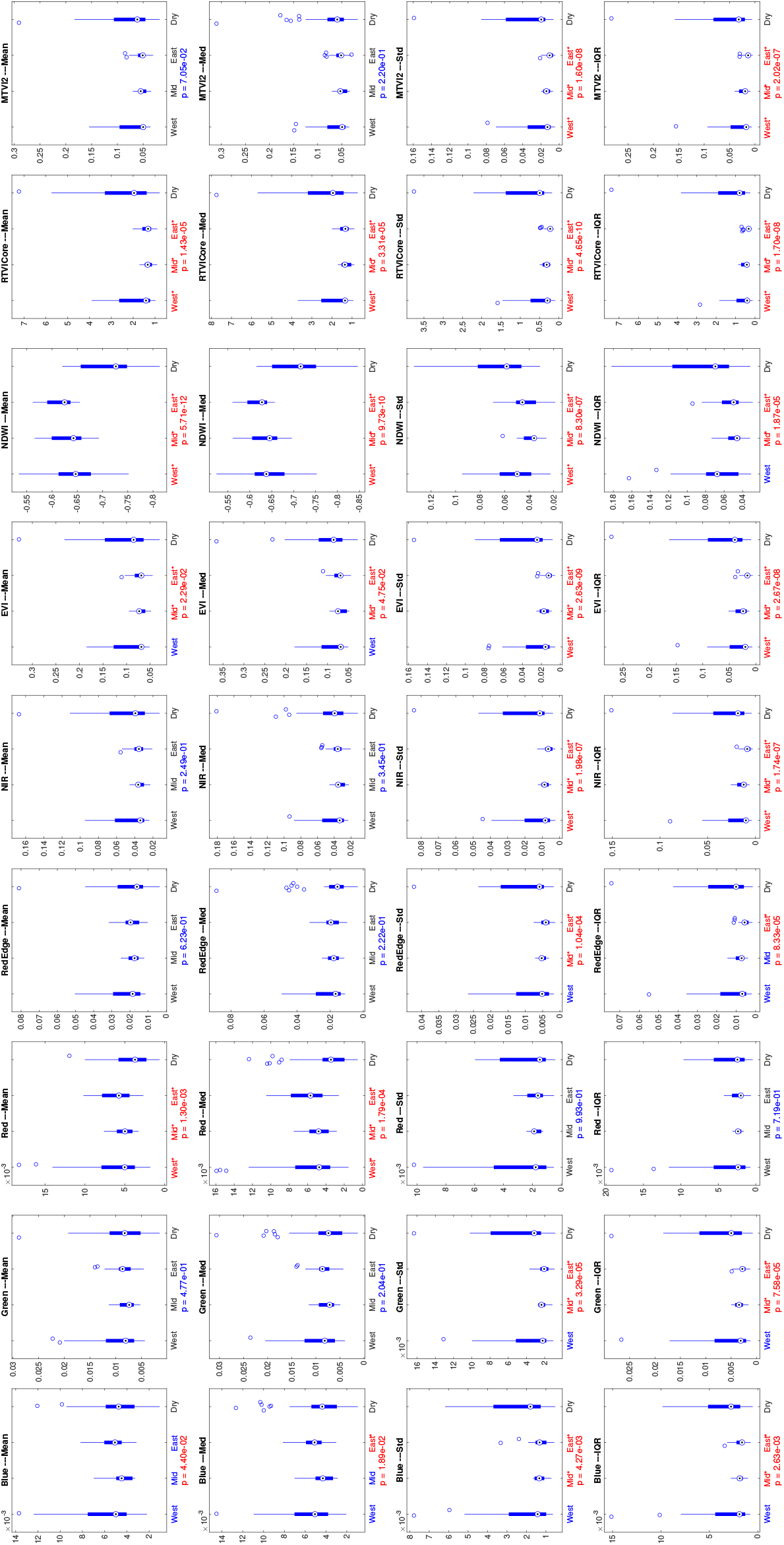
Boxplots indicating discrepancies between the local measurements in each of the four studied areas (western, middle and eastern parts of the peat bog, with continuously reduced elevation, as well as dry land areas at surrounding elevations), including five multispectral bands (Blue, Green, Red, RedEdge, NIR), as well as calculated vegetation indices (EVI, NDWI, RTVICore, MTVI2). Statistically significant discrepancies are indicated by red color, with annotated *p*-values according to one-way non-parametric ANOVA (Kruskal-Wallis) statistical test, particular areas that are different from dry land (acting as control) are denoted by asterisks.

